# Signal Attrition in Whole Cell Crosslinking Mass Spectrometry

**DOI:** 10.1101/2025.10.16.682930

**Authors:** Bruno C. Amaral, Nicholas I. Brodie, Andrew R.M. Michael, D. Alex Crowder, Pauline Douglas, Morgan F. Khan, David C. Schriemer

## Abstract

Crosslinking mass spectrometry (XL-MS) could replace traditional techniques for sampling the cellular interactome, such as affinity pulldown MS. It generates superior interaction data that can be used to better model cellular structure and function. However, the sampling depth of current XL-MS techniques is disappointing. Poor sampling is often blamed on a low-yielding crosslinking reaction, estimated to be ∼0.1% based on relative ion abundance in the mass spectrometer. Here, using a new two-step crosslinker installation process, we demonstrate that the yield of the chemical reaction is not the limiting factor in sampling the interactome. Low crosslink detection levels persist even when crosslink yields approach 30% of total peptide. We show that crosslinked peptides are not preferentially lost during sample workup; they enter the mass spectrometer at least as efficiently as linear peptides. Low detection rates arise mostly from severe signal splitting that reduces the S/N of crosslinked peptides, which is likely exacerbated by low-sensitivity database search tools and compounded by ion suppression.

## Introduction

Crosslinking mass spectrometry (XL-MS) is an emerging method in structural biology and interactomics^1^. Protein crosslinks are installed through the application of bifunctional chemical reagents, which are then measured by MS using well-established bottom-up proteomics methods. XL-MS is highly complementary to modern structure prediction methods because it can validate structures and even enhance model accuracy^2–4^. When applied to cells, the experiment generates distance measurements that can positively identify a protein-protein interaction (PPI) and even support *in situ* structure determination^5,6^. It could become a preferred method for mapping the organization and flux in the cellular interactome, as all other techniques for interaction mapping return only indirect information.

Major advancements in whole-cell crosslinking have been made, but the depth of interactome sampling remains low^7–9^. Even extensive efforts struggle to generate more than hundreds of PPI’s per experiment. One reason often given for poor performance is the yield of the desired reaction product: the crosslinked peptide. It is estimated to be exceptionally low.

About 0.1% of the MS signal can be attributed to crosslinked peptides and there appears to be a bias towards the highly abundant proteins in complex samples^8,10,11^. These observations have fostered an analytical strategy that incorporates enrichable crosslinkers and extensive fractionation of proteolyzed cell lysate^6,12–14^. While such methods have improved interactome sampling, the numbers remain low and far removed from a full sampling of the predicted complexity of the interactome^15^. The only viable way to improve coverage seems to involve large amounts of cell lysate as input combined with extensive multimodal enrichment technologies, which makes the technique impractical as a tool for routine biological inquiry.

In this study, we set out to explore the reasons for low performance in greater depth, taking advantage of a new method of crosslink installation that overcomes several weaknesses in the classical approach (**Figure 1**). Here, the cell is first rapidly stabilized using high-concentration formaldehyde (4% v/v), rendering the protein contents of the cell immobile yet accessible to a multi-step crosslinking process^16^. Two crosslink precursors are then installed on surface-accessible lysines, followed by a catalyzed click reaction to achieve the desired crosslinks. The benefits of this “click-linking” approach are numerous. First, formaldehyde crosslinking does not compete with amine-targeting crosslinkers, if formaldehyde treatment is limited to the conventional 10-15 min used in immunofluorescence experiments. Second, the spatial proteome is not strongly distorted during the reactions^17^. Fixation using the standard 4% formaldehyde solution preserves the location of most proteins in the cell with high fidelity, although there can be some artefacts^18^. Third, conventional crosslinking of cells and lysates generates a bias towards high-abundance proteins^11^. Prefixing the cell limits the diffusion (or mixing) of proteins upon which this phenomenon is based. Fourth, the crosslinking step is reduced to a single coupling, with no competition arising from reagent hydrolysis, other biomolecules, or even with itself^19^. Fifth, because the precursor installation step is independent of the actual crosslinking reaction, the method should offer control over crosslinker yield, simply by varying the level of click-linking precursors that are installed in the first step. Here, we show that click-linking can indeed generate very high levels of crosslink products. These do enter the mass spectrometer, but the efficiency of detection remains extremely low primarily due to signal splitting.

**Figure 1.**
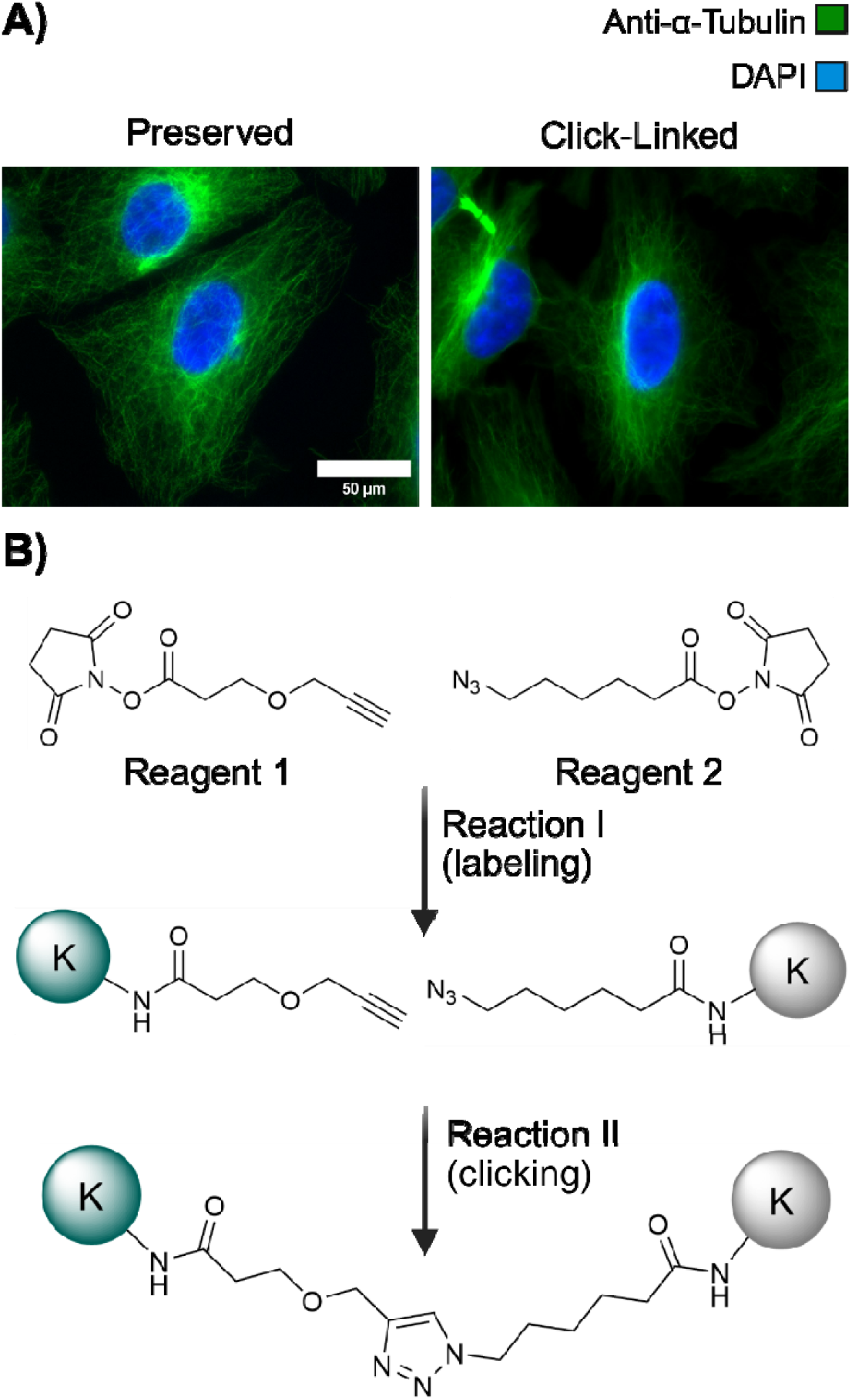
The click-linking procedure. (A) Cells are first fixed and stabilized with high concentrations of formaldehyde, as in a typical immunofluorescence microscopy experiment (left image, preserved). After fixation and washing, NHS-activated reagents compatible with a copper-catalyzed azide-alkyne cycloaddition (CuAAC) reaction are installed in a 1:1 molar ratio, followed by a second wash and the addition of Cu(I) and an accelerating ligand to complete the crosslinking reaction (right image, click-linked). Click-linking causes no obvious distortion of cellular ultrastructure. α/β-tubulin shown in green (using FITC-labeled anti-α-tubulin) and nucleus shown in blue (using DAPI stain). (B) Click-linking reagents used in the current study, 2,5-dioxopyrrolidin-1-yl 3-(prop-2-ynyloxy)propanoate (**1)** and (2,5-dioxopyrrolidin-1-yl) 6-azidohexanoate (**2**, showing the two-step nature of the crosslinking reaction.

## Methods

### Click-linking protocol

A549 cells were grown to 80% confluency in Ham’s F-12K medium (Gibco) supplemented with 10% fetal bovine serum (FBS, Gibco) and 1% penicillin-streptomycin (Gibco) at 37 °C in a humidified atmosphere with 5% CO_2_ on 10 cm plates. Cells were washed 3x with PBS to remove media and then incubated for 10 minutes in 5 mL of a solution of PBS and 4% formaldehyde. After formaldehyde treatment, cells were washed 3x with PBS and treated with 5 mL of 0.1 % Triton in PBS for 10 minutes to permeabilize the cells, followed by 3x washes with PBS. Next, cells were treated with 1 mM each of 2,5-dioxopyrrolidin-1-yl 3-(prop-2-ynyloxy)propanoate (**1**) and (2,5-dioxopyrrolidin-1-yl) 6-azidohexanoate (**2**), each diluted from a 100 mM stock with PBS and added simultaneously (5 mL total, incubated for 1 hour at 25°C with gentle agitation). Cells were washed 3x with PBS and incubated for 5 minutes between each wash. For the 3x labeling experiment, this procedure was repeated twice. The click reactions were performed by adding, in order, 5 mM copper chloride, 10 mm BTTES and 50 mM sodium ascorbate, in a total volume of 5 mL of PBS. Click reactions were incubated for 30 minutes at 25°C, then washed 3x with PBS for 5 minutes each. 11,12-didehydro-γ-oxodibenz[b,f]azocine-5(6H)-butanoic acid (DBCO-acid) was added to a final concentration of 2 mM in 5 mL to cap remaining azides. Three final washes with PBS were performed, and the cells were then lysed and recovered in detergent-containing lysis buffer containing 500 mM Tris, 150 mM NaCl, 1x protease inhibitor, 1% Triton, 2% SDS, 6 M Urea and 5 mM DTT and collected in a microfuge tube. Where appropriate, a detergent-free lysis was performed by scraping cells into 500 µL of PBS and lysing on ice using a Fisher Scientific sonic dismembrator 100 at power setting 5, producing approximately 9 W output power, for 13 seconds (3 seconds on, 10 off, 4 cycles).

### SP3 processing of cell lysates for MS analysis

Lysate (with or without detergent as appropriate) was treated with 6 M Urea and 10 mM DTT for 30 minutes. 2-chloroacetamide (CAA) was then added to a final concentration of 40 mM and incubated for 40 minutes at room temperature. An equal mixture of the two SP3 bead slurries was prewashed and added to protein sample in a 1:10 (v/w) ratio of bead slurry to protein mass, and ethanol was added to 50% (v/v) to precipitate proteins on the beads. Samples were incubated for 15 minutes with agitation at room temperature. Beads were washed with 80% ethanol three times, transferring beads to a new tube after the second wash. Samples were digested in 50 mM ammonium bicarbonate (AB) pH 7.8 with trypsin in a 1:50 (w/w) enzyme:protein ratio overnight at 37°C and then dried by speed vac. Samples were resuspended in 20 µL of 0.1% TFA and desalted by ZipTip following the manufacturer’s standard procedures (Millipore Sigma, ZTC18S960). Peptides were eluted using 80% ACN/0.1% FA, dried, then resuspended in 20 µL of 0.1% FA and approximately 1 µg was injected into the LC-MS.

### Preparation of supernatant and pellet for comparative analysis of digests

100 µL of lysate (in detergent-containing lysis buffer) was pelleted at 10,000 x g for 15 minutes. The supernatant was removed, and the pellet was washed once with 100 µL of PBS, followed by resuspension in 100 µL of PBS. Both total supernatant and pellet were treated to SP3 processing as above, and 2 µg of trypsin was used for overnight digestion at 37°C. Resulting digests were dried by speed vac and resuspended twice to remove AB and then finally resuspended in 20 µL of 0.1% FA for injection on the mass spectrometer. Equal volumes were injected for a direct comparison of intensity.

### Preparation of cells for amino acid analysis

A549 cells were click-linked as above, where two populations were generated: pre-clicked (where only the precursors were installed) and post-clicked (after crosslink formation). Control A549 cells were also prepared that were simply fixed and permeabilized, then washed with PBS. Cells were harvested with 1 mL of lysis buffer, excluding DTT. 1 mL of lysate was spun down at 10,000 x g for 10 minutes (4°C). For each of the three samples, the pellet was washed a total of three times with PBS, spinning after each wash. Pellets were snap frozen and shipped for amino acid analysis on dry ice, performed at the SPARC BioCentre (Sick Kids Hospital, Toronto) using standard protocols.

### Colorimetric peptide assay

Digests were prepared as above. The resulting dried product was resuspended in 30 µL of 25 mM AB, and 10 µL of the product was analyzed using the Pierce colorimetric peptide quantification kit (product #23275, Thermo Scientific), as per the manufacturer’s protocol. Briefly, 95 µL of the working solution was added to the 10 µL of sample. A 5 point standard curve was prepared (ranging from 0.0313-0.5 mg/mL) from the stock peptides in the kit. Absorption measurements were taken in a 384 well plate, using a Molecular Devices spectrophotometer (FilterMax F5) at a wavelength of 450 nm. For the analysis of the ZipTip-purified peptides, 4 µg of peptides, recovered after SP3 and quantified, was acidified and ZipTipped as above. Recovered peptides were dried in the speedvac and resuspended in 10 µL of 25 mM AB for colorimetric quantitation, as above.

### Fractionation

Samples (either pre-clicked or post-clicked) were fractionated using a Vanquish Online 2D-LC system (Thermo Fisher Scientific). For high-pH fractionation, desalted peptide samples were resuspended in high-pH Mobile Phase A (20 mM ammonium formate, pH 10) and loaded onto a ZORBAX RRHD Extended-C_18_ column (80Å pore size, 2.1 x 150 mm, 1.8 μm particles, Agilent). Separation was performed in a multistep 68 min gradient at a 200 µL/min flow rate, from fixed 5% B (80% acetonitrile) for 1 min; 5-40% B for 42 min; 40-60% for 8 min;60-95% for 2 min; fixed 95% for 4 min; then ramped back 95-5% B for 2 min, holding at 5% for another 10 min. Eluate fractions were collected every minute from 1-61 min for a total 60 fractions, which were concatenated into 30 final fractions (fraction 1+30, 2+31, etc), dried down and resuspended in 0.1% formic acid for LC-MS/MS analysis. For the clicked samples that were additionally fractionated by size-exclusion chromatography (SEC), peptides were resuspended in SEC Mobile Phase (0.1% formic acid, 30% acetonitrile) and loaded onto a Superdex peptide PC 3.2/30 column (GE Healthcare). Separation was performed in 60 min using an isocratic flow of Mobile Phase at 0.08 mL/min. After 10 min, eluate fractions were collected every 1.9 min for a total of 24 fractions. Out of the first 11 fractions, the first 7 were concatenated into 1, and fractions 8, 9, 10 and 11 were kept individually, while fractions 12 through 24 were discarded.

Each of the 5 final fractions were submitted to the high pH fractionation and collection method described above. A total of 150 samples were dried and resuspended in 0.1% formic acid for LC-MS/MS analysis.

### LC-MS/MS

Data were collected using a Vanquish Neo UHPLC coupled to an Orbitrap Ascend mass spectrometer fronted by an Easy-Spray Source (ThermoFisher Scientific). For each sample, approximately 1 µg of peptide mass was injected onto a 300 µm x 5 mm PepMap Neo Trap Cartridge peptide trap column (C18, 5 µm particle size, 100 Å pore size, Thermo Fisher Scientific), which were then submitted to reverse phase chromatography using an EASY-Spray HPLC analytical column (75 µm x 50 cm, C18, 2 µm particle size, 100 Å pore size, Thermo Fisher Scientific) at a flow rate of 250 nL/min at 40°C. Mobile Phase A consisted of 0.1% (v/v)formic acid in water, while Mobile Phase B consisted of 0.1% (v/v) formic acid in 80:20 acetonitrile:water (v/v).

For precursor-labeled samples, separation was carried using a multistep 120 min gradient from 4-30% B for 85 min; 30-45% B for 20 min; then a 15 min wash at 99% B. MS data were acquired in positive mode using data-dependent acquisition (DDA) with a 2 second cycle time. Full MS scans were performed in the orbitrap set at 120,000 resolution (*m/z* 200) for the 375-1875 *m/z* range, with maximum injection time set as Auto and normalized AGC target as Standard. Precursors with charge states 2-6+ were selectively isolated for fragmentation using stepped HCD at normalized Collision Energies (nCE) of 27, 30 and 33 (quadrupole isolation window of 1.6 Th). The Dynamic Exclusion window was set to 10 seconds. MS^2^ scans were performed using the orbitrap set at 30,000 resolution (*m/z* 200) with maximum injection time set to Auto and the AGC target to 1e5.

For clicked samples coming from fractionation, separation was carried out using a multistep 90 min gradient from 4-20% B for 55 min; 20-45% B for 15 min; then a 20 min wash at 99% B. MS data were acquired using DDA in a 2.5 sec cycle time using mostly the same parameters, with some modifications. Full MS scans were performed in the orbitrap at 120,000 resolution (*m/z* 200) for the 375-1875 *m/z* range, with maximum injection time set to Auto and AGC target at 1e6. Precursors with charge states 4-8+ were selectively isolated for fragmentation using stepped HCD at Normalized Collision Energies (NCE) of 27, 30 and 33 (quadrupole isolation window of 1.6 Th). The Dynamic Exclusion window was set to 10 seconds. MS^2^ scans were performed using the orbitrap set at 30,000 resolution (*m/z* 200) with maximum injection time set to Auto and AGC target to 1e5.

For samples used to investigate possible sample loss, we used a gradient of 2%-30% B over 3 hours, and 30-45% over 1 hour. MS spectra were acquired at a 200% AGC target at 120,000 resolution in the Orbitrap. MS^2^ spectra were collected at 300% AGC at 30,000 resolution in the Orbitrap. Either +2-8 charge states were acquired, or +4-8.

### Data Analysis

MSFragger^20^ analysis using the default LFQ-MBR search routine was performed against the full UniProt Human proteome database with the following parameters: cysteine carbamidomethylation (Δm = +57.021465 Da) as fixed modification; methionine oxidation (Δm = +15.9949 Da) and N-termini acetylation (Δm = +42.0106 Da) as variable modifications; 10 ppm for precursor mass tolerance; 10 ppm for fragment mass tolerance; trypsin as enzyme for protein digestion with 3 allowed missed cleavages. The mass shifts corresponding to the reagent modification were adjusted as follows: Reagent **1** (Δm = 110.0366 Da), Reagent **2** (Δm = 139.0745 Da), DBCO-Capped-Azide (Δm = 444.1798 Da). IonQuant was used for label-free quantitation and each mass spec run was processed individually. The LFQ-MBR workflow was used to investigate the quantitative proportion of labeled versus unlabeled peptides and estimate reaction performance according to the following equation:

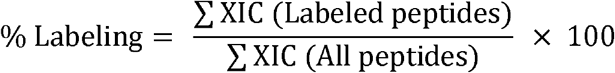

Crosslinks were identified using multiple tools. When using pLink 3.0.17^21^, the parameters used were as follows: peptide length 6-60; trypsin as the digestion enzyme with 3 allowed missed cleavages; carbamidomethylation of cysteines (Δm = +57.021 Da) as a fixed modification; oxidation of methionine (Δm = +15.995 Da) as a variable modification; 5 ppm for precursor and 10 ppm for fragment mass tolerance. Data were searched against the full UniProt Human proteome databases at 5% false discovery rate (FDR) at the spectrum level. Crosslink (Δm = +249.111 Da) and monolink modifications were added for the clicked products, with specificity to Lysine residues and protein N-termini. Charge states were analyzed in Freestyle, which outputs the precursor peptide charge for all MS^2^ spectra acquired. For trend comparisons, other crosslink search tools were used. For pLink v2.3.11, the same parameters as the newer version were used. For MS Annika 3.0, default parameters provided in the DSS/BS3 MS^2^ template for large datasets were used, with a few modifications: 5ppm for MS1 tolerance, 10ppm for MS^2^ tolerance;); peptide length 5-30; TopN Filter 5; static modification of cysteine carbamidomethylation (Δm = +57.021 Da); dynamic modifications of methionine oxidation (Δm = +15.995 Da) as well as the respective crosslink and monolinks masses. For CRIMP 2.0, default parameters were used with the addition of: carbamidomethylation of cysteines (Δm = +57.021 Da) as a fixed modification; oxidation of methionine (Δm = +15.995 Da) as a variable modification; 5 ppm for precursor and 10 ppm for fragment mass tolerance, minimum fragment threshold = 3, maximum and minimum isolation window = 0.8, %E-Threshold = 50%.

To estimate the number of linkable sites per lysine, a random selection of 110 PDB files were analyzed in Jwalk^22^ (version obtained from GitHub) using default parameters that limit linkages to a surface accessible distance less than 30Å. From these results, python scripts were used to scrape the number of linkages and the number of lysines from each of the PDB files, and the average number of linkages per lysine for each structure was calculated simply by dividing the number of linkages by the total number of lysines in the structure. Estimates of crosslink abundance were made by generating extracted ion chromatograms (XICs) for all detected features in matched datasets. That is, identifications of all linear peptides from a long-gradient shotgun data acquired by triggering MS^2^ on 2+ and 3+ ions were transposed on a matched sample acquired by triggering on 4+ and higher, using match-between-runs concepts. This allowed for XICs to be measured for all reaction products in one run, and relative crosslink abundance determined simply from a ratio of total crosslink XICs to all XICs.

## Results and Discussion

To investigate the actual yield of the crosslinking reaction, we first applied the click-linking protocol to human A549 cells and measured the detectable click-link precursors before and after the click reaction. Briefly, we treated pre-stabilized A549 cells with a single application of an equimolar mixture of 2,5-dioxopyrrolidin-1-yl 3-(prop-2-ynyloxy)propanoate (reagent **1**) and (2,5-dioxopyrrolidin-1-yl) 6-azidohexanoate (reagent **2**) (**Figure 1**). Then, instead of clicking, we harvested and processed the cell lysate for a shotgun proteomics experiment, to determine the level of labeling via measured peptide intensities. In the matched sample, we labeled the A549 cells as above and then initiated the click-linking process by adding the catalysts and allowing the reaction to proceed to completion. The most effective way to estimate the yield of crosslinking is to measure the difference in precursor abundance between these two states. It is preferred over the direct detection of crosslinked peptides because the same species is tracked in both samples (the precursors) rather than a new form (the crosslinked peptide), thus avoiding concerns over variable ionization efficiency. The calculation is based upon the measured intensity of the identified peptides (see methods). Mono-labeled peptides can be detected with high efficiency and because the click reaction is highly selective, any reduction in the observed fractional levels of precursors can only arise through the generation of protein crosslinks^17^. This approach was corroborated by fluorescence measurements before and after crosslinking^17^, and we also note that side reactions with the click reagents were not detected (**Figure S1**), in keeping with the bio-orthogonality of the click reagents.

We then installed click-linking reagents over a range of concentrations, from 0.1 mM to 3 mM per reagent, and used long-gradient shotgun runs to measure precursor levels. Note that the 3 mM application refers to three equal and consecutive 1 mM reactions, with washing steps in between, as we are limited by the solubility of the reagents. Such an approach is possible because the cells are fixed and stabilized, as noted above. Labeling yields increased steadily, up to 43.6% of total signal at 3x1 mM, and the levels of detectable precursor dropped dramatically after click-linking for all concentrations (**Figure 2A**). For the 1 mM reaction, 35.3% of all peptides measured were labeled, which dropped to 7.9% after click-linking. The level of installed precursors never goes to zero regardless of the length of the reaction (not shown). Although the reagents are installed in a 1:1 ratio, alkyne-to-alkyne and azide-to-azide mismatches can occur and render some fraction of lysines unavailable for crosslinking. Additionally, some precursors will not be proximal to other precursors, given the combined length of the two reagents (∼18Å), or because the reagents are not equally dispersed throughout the proteome. Further topological incompatibility may also arise from a small fraction of the precursors that penetrate protein structure and label (partially) buried lysines. Nevertheless, 27.4% of the total signal is converted into protein crosslinks in the 1 mM reaction, which greatly exceeds the 0.15% estimated from measured crosslink ion abundances in a conventional PhoX crosslinking experiment^10^. Fluorescence measurements have verified this high yield^17^, as we have noted above.

**Figure 2.**
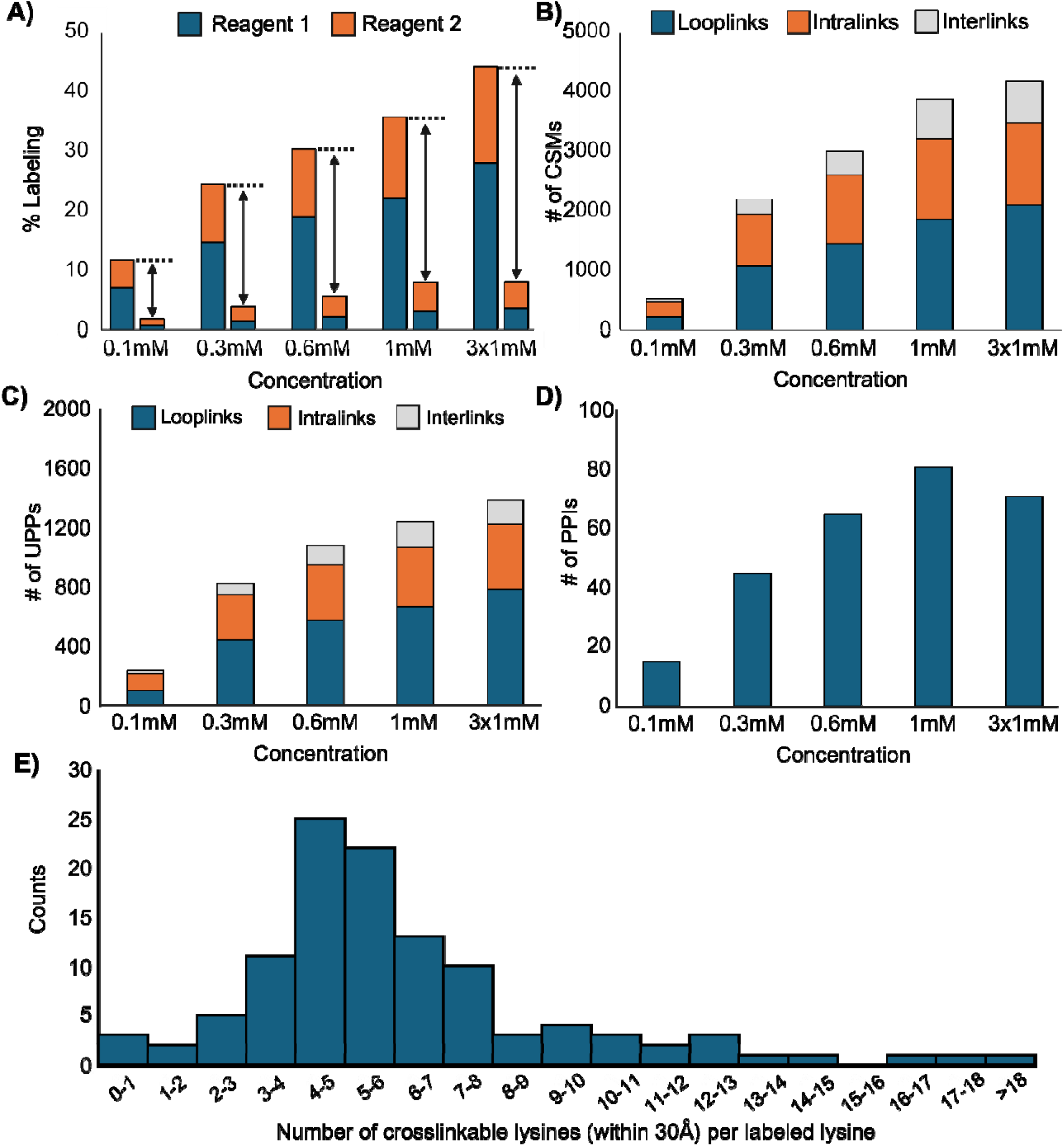
Maximizing crosslink yield in the human proteome. (A) Detected levels of click precursors installed in the proteome, and the reduction in their levels upon click-linking, across the indicated concentration range for reagents **1** and **2**. (B) CSMs (C) UPPs and (D) PPIs detected from targeted acquisitions of crosslinks across the same concentration range. (E) Calculation of the average number of spatially accessible crosslink points per lysine residue, using Jwalk and based on a random sampling of protein structures.

We then conducted long-gradient shotgun runs for each click-linked sample in the concentration series and biased the ion selection towards crosslinks by targeting higher charge states (+4-8). The level of crosslinked spectrum matches (CSMs) and unique peptide pairs (UPPs) increased with reagent concentration, but not proportionally, and the number of detectable PPIs actually dropped slightly in the 3x1 mM reaction **(Figure 2B-D**). When we conducted an XL-MS analysis of the 1 mM reaction at greater depth, using milligram amounts of input material and extensive fractionation (SEC and 2D RP-LC), the experiment netted 28,462 unique crosslinked peptides and an estimated 600 - 1700 PPIs, depending on the database search tool that we used^17^. The combined intensity of the crosslinked peptides that we detected was 0.3% of the total signal (see methods). This low yield of detected crosslinks is only somewhat better than previous estimates, and still inconsistent with the high yield of the reaction that we calculated from the precursor detection approach.

To investigate the discrepancy between these two different measurements of yield, we first calculated how many crosslinks should be introduced into the mass spectrometer in a click-linking experiment, based upon some reasonable approximations. First, we conducted an analysis of structural knowledgebases to estimate how many crosslinks might be created for every surface-accessible lysine. We assume that distances up to 30 Å between α-carbons of proximal lysines can be linked using a crosslinker of this length (∼18Å). We sampled protein structures until the nature of the distribution became apparent and observed that, on average, one lysine can couple with six other lysines (**Figure 2E**). The protein databank (PDB) that we sampled is mostly populated with structures of individual proteins, but if we assume that intraprotein crosslinks dominate our crosslinks^23^, then the calculation represents a reasonable lower limit of crosslink complexity.

We then measured how many lysines could be labeled in the first step of the click-linking reaction, using a deep-sampling 2D LC fractionation scheme like that used in the XL-MS workflow. We detected 137,337 lysine-containing peptides. This set reduced to 77,200 unique lysines, of which 50,900 (66%) were labeled with either or both precursors (**Figure S2**). To estimate how many crosslinked peptides might be generated from a pool of this size, we note that essentially all lysine-containing peptides either disappear or drop in relative intensity by 50% or more after the click reaction (**Figure S3**), so we will assume that all participate in one or more crosslinks. For simplicity, we will approximate the linkable lysines in a spatial proteome as an infinite 2D hexagonal array with 30 Å spacing between lattice points. This allows us to estimate the number of crosslinks formed to be six linkages times 50,900 lysines, divided by two to avoid double-counting, for a total of 152,700 unique lysine-lysine crosslinks. This number very likely represents a lower bound on the crosslinks formed. This calculation only accounts for precursor peptides in a nominally MS-detectable range. The whole proteome contains 650,000 lysines, and following a similar calculation would yield 1.95 million potential unique crosslinks. Our actual direct measurement of the 1 mM click-linking experiment (28,462 unique crosslinks) is clearly well below both estimates.

We then wondered if sample handling introduced a bias against crosslink peptide delivery to the mass spectrometer. We noticed upon workup of the crosslinked cells that the lysate contained precipitate at levels that appeared to correlate with the degree of click-linking: increasing the level of crosslink precursor increased the cloudiness of the lysate. To determine how much protein was in the precipitate, we centrifuged and washed it extensively to remove soluble protein, and then conducted quantitative amino acid analysis on the pellet^24^. This analysis incorporates a harsh acid hydrolysis, ensuring that any protein in the precipitate would be resolubilized for quantitation. Three samples were prepared and analyzed: pelleted lysate from control cells that were not treated, pelleted lysate from cells treated only with 1 mM of the click precursors, and pelleted lysate from cells subjected to the full click-linking process (**Figure 3A**). The amount of protein in the click-linked sample pellet was 2.5 times greater than the protein in the pellet of the control. The lysis buffer we used did not appear sufficient to solubilize very much crosslinked protein. The supernatant was essentially free of crosslinks (**Figure 3B**).

**Figure 3.**
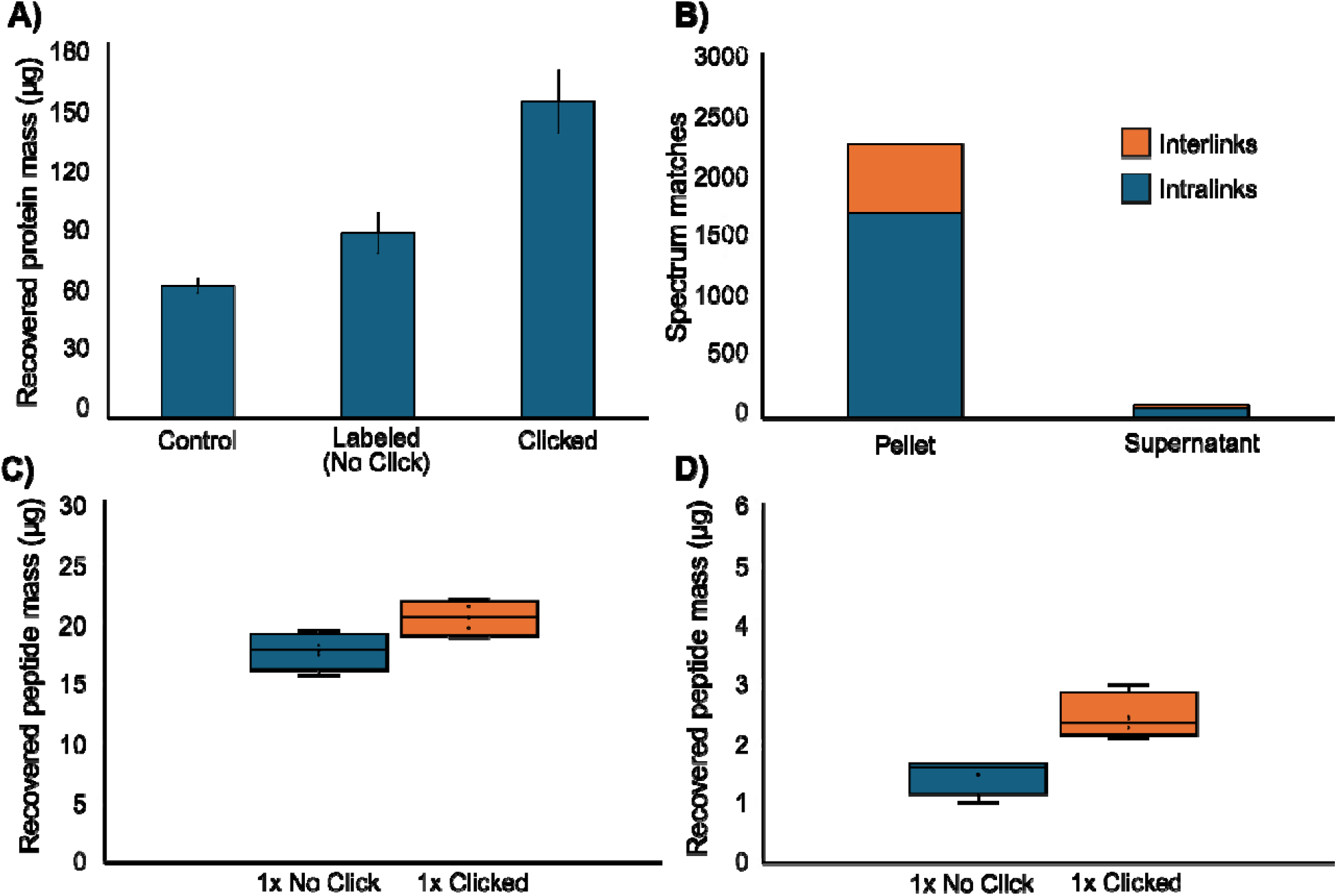
Crosslink peptide recovery after click-linking. (A) Increased protein precipitation upon click-linking as quantified by amino acid analysis. Error bars represent 1 std. dev. (n=3 acquisitions). (B) Crosslink spectrum matches detected via shotgun proteomics analysis of the digested pellet and the supernatant. (C) Quantification of peptide recovered from SP3 beads using a colorimetric peptide assay. Error bars represent 1 std. dev. (n=3 acquisitions). (D) Quantification of peptide recovery from a C18 zip-tip, using the same colorimetric assay. Error bars represent 1 std. dev. (n=3 acquisitions).

Extensive crosslinking leading to precipitative loss could provide ample reason for a lower detection rate, but most of the pellet was solubilized when trypsin was added. We explored several different lysis buffer formulations and discovered that a variant of RIPA buffer (50 mM Tris pH 7.4, 150 mM NaCl, 0.5% NP-40, 1mM EDTA) prevented protein precipitation after click-linking, although it resulted in virtually identical cross-link yields. Thus, precipitative loss does not appear to be a major reason for the low rate of crosslink detection.

We then tested if crosslinking impeded digestion, or our ability to recover crosslinked peptides and deliver them to the mass spectrometer. We examined the efficiency of peptide recovery using MS analyses and a colorimetric peptide assay and first investigated digestion performance. Our digestion protocol incorporates SP3 technology that is used extensively in clinical proteomics, involving bead-based capture of protein from lysates followed by on-bead tryptic digestion^25^. The method is effective at removing buffer components that would otherwise interfere with downstream LC-MS analysis, while promoting good recovery of digested material. It uses an organic solvent to aggregate and disperse protein on carboxylate-modified magnetic beads, rendering the sample easy to wash and recover after digestion^26^. Interestingly, the SP3 method appeared to be more efficient at recovering peptide from click-linked samples than the corresponding unclicked control (**Figure 3C**), and an exploration of the search results showed that linear peptides were detected with greater efficiency after click-linking (*i*.*e*, over 50% more linear peptides). The click-linked sample is already prone to precipitating, so it seems sensible that total protein recoveries could be higher than unclicked samples and thus increase total peptide recovery. Indeed, a variant of SP3 forgoes the beads altogether in favor of direct solvent-induced precipitation, where recoveries can be superior to SP3^27^.

It is also possible that crosslinking rendered the crosslinked peptides more difficult to release from the SP3 beads. To investigate, we digested the bead-bound protein in the presence of elevated ionic strength (to weaken the interaction with the carboxylate-modified beads), and separately with 10% acetonitrile (to weaken hydrophobic interactions), but neither changed the distribution of reaction products. Additional peptide was recovered from the beads with a second elution protocol involving 10% FA, but again, the distribution of reaction products was not measurably different (**Table S1**). Finally, we also tested if click-linking prevented protein from binding to the SP3 beads, but we detected nothing in the supernatant. In sum, biased recovery does not contribute to the low rate of crosslink detection, and total peptide recoveries are actually superior to the control.

We then evaluated the effect that sample clean-up might have in generating a bias against crosslinked peptides. We incorporated a C18-based Zip-Tip for final sample polishing after recovery from the SP3 beads. We applied a matched amount of total unclicked and clicked digest to the Zip-Tips (4 µg), and then tested the eluate from the tips using the colorimetric assay. We found that, here as well, peptide recovery from the clicked sample was superior to the unclicked control, suggesting no preferential loss in C18-based purification and separation (**Figure 3D**).

Indeed, it indicates that crosslinked peptides are recovered somewhat more efficiently than linear peptides. Taken together, these results suggest that crosslinked peptides are produced in high amounts and delivered to the LC-MS system with an efficiency meeting or exceeding that of linear peptides.

We then evaluated MS-related parameters. We first investigated if the crosslinks themselves are unstable during ion transmission, which would obviously reduce detection efficiency. But in-source fragmentation is essentially non-existent, an observation that is consistent with the stability of the aromatic triazole linkage. Only the most labile of crosslinkers can be induced to undergo extensive fragmentation in the source^28^. These findings are consistent with the absence of gas-phase cleavage in the MS^2^ spectra of identified click-linked peptides (**Figure S4**). Thus, in-source cleavage does not account for the discrepancy we observe between crosslink yield and the detected crosslinked peptides.

Lastly, we examined the spectral output from the DDA experiments, and the conversion of precursor selections into CSMs. XL-MS experiments usually select higher charge states than a typical bottom-up proteomics experiment, following the logic that crosslinked peptides are larger and thus would have a charge state distribution shifted to higher values. We examined ion sampling as a function of charge state, by letting the instrument select precursors in two ways (**Figure 4A**). We first sampled within the +2-8 range for both the clicked and unclicked samples and then narrowed the range to +4-8. For the former, well over 90% of the precursor ions were +2 and +3 charge states, regardless of sample type. When focusing exclusively on 4-8+ ions, there was an overall 5-fold increase in the number of precursor ions selected for MS^2^ for this charge state range (13,361 to 71,689) (**Figure 4B**) but this number is still quite small compared to the number of crosslinks created with the click-linking protocol. We then examined the spectral conversion efficiency for the 4-8+ DDA experiment, that is, the success rate in converting a selected precursor ion into a CSM, scored at a nominal 5% FDR. The conversion efficiencies steadily decline as a function of charge state, dropping to near zero for 8+ precursors (**Figure 4C,D**). Interestingly, although there is a slightly higher conversion efficiency upon click linking, this metric sums both linear and crosslinked peptides. The 5-fold increase in high charge-state sampling in the +4-8 acquisition is mostly linear peptides; only 1% of this boost resulted in new CSMs. In the aggregate, these analyses suggest that crosslinked peptides are not being sampled in high numbers because their intensities are very low, with perhaps an additional penalty arising from low sensitivity CSM scoring functions.

**Figure 4.**
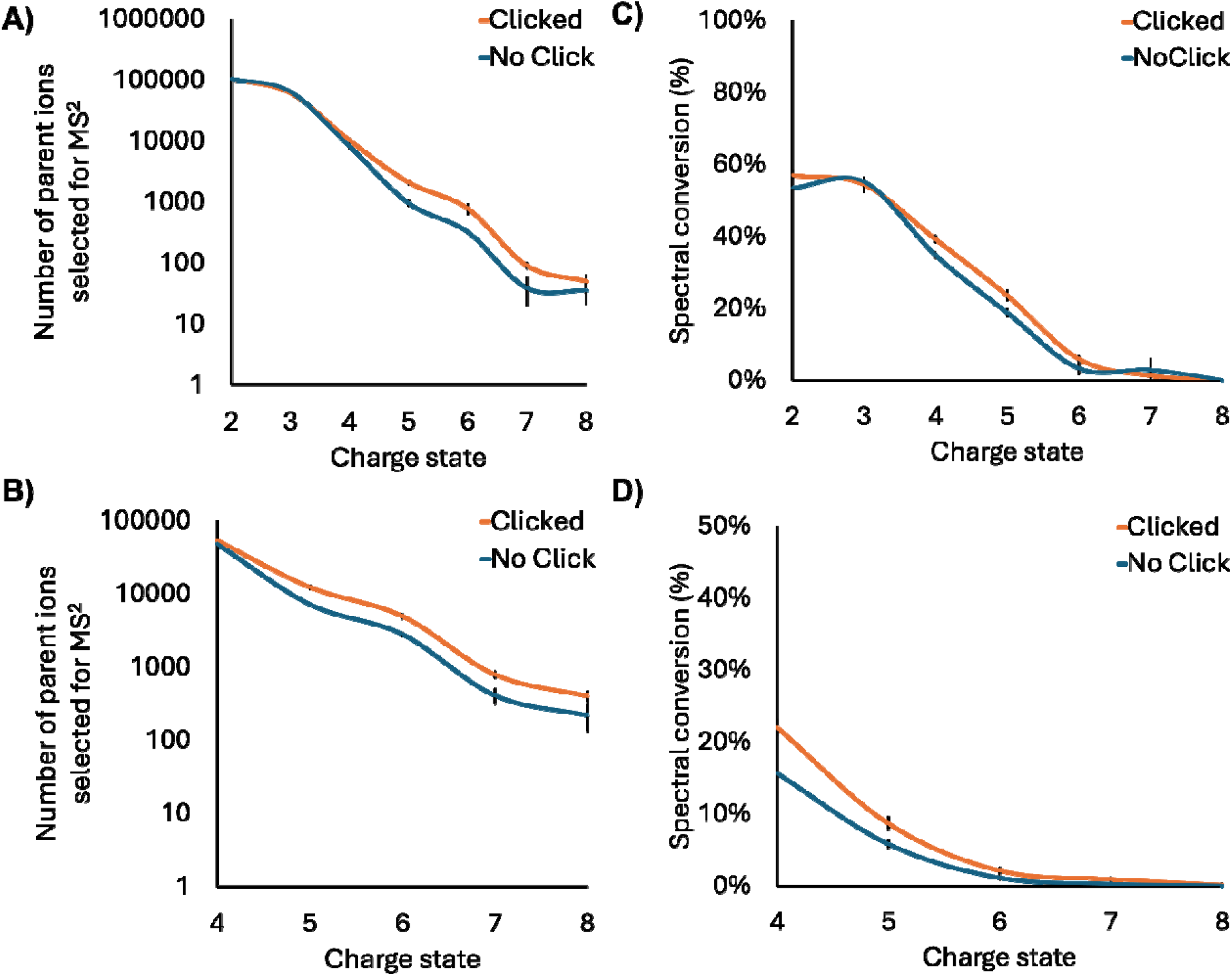
Effect of click-linking on ion selection and identification. (A) Number of ion selected for MS2 analysis as a function of charge state for both the click-linked samples and the non-clicked control samples, for acquisitions targeting charge states +2-8. (B) As in A, but for acquisitions targeting charge states +4-8. (C) The percentage conversion of selected ions into hits above the FDR-controlled score cutoff, for acquisitions targeting charge states +2-8. (D) As in C, but for acquisitions targeting charge states +4-8. Error bars in all graphs represent 1 std. dev. (n=4 acquisitions).

### Assessment

Our study conservatively estimates that at least 150,000 lysine crosslinks are created in a click-linking experiment (and likely much more), and the linked peptides are efficiently transferred to the mass spectrometer. Why are they not detected? One reason could involve the restrictions that we apply to the database search. We typically restrain our search to peptides with greater than six residues to ensure accuracy in the PPIs returned. It means that only ∼70% of the peptides in an “Arg-C” digest can be converted into discriminating hits^29^. (Arg-C is an appropriate cleavage rule if we assume that all surface lysines are labeled and undigestible by trypsin, see **Figure S2**). This restriction could reduce our estimate of detectable crosslinked lysines to. We observed 28,818 unique crosslinked peptides, but this collapses to only 7492 unique lysines, as our detected crosslinks are represented by multiple overlapping peptide sequences.

Thus, only 10% of our most conservative estimate of detectable crosslinked lysines are observed, and far less than the actual lysines in the proteome. We primarily used pLink3 in this study. The number of detectable crosslinks is somewhat dependent on the search tool (**Figure S5**), but pLink3 generates the highest number by a considerable amount (consistent with an assessment of the previous version as a rather permissive tool^30^). Thus, it is possible that we are even overestimating the number of detectable crosslinks. As noted above, our database search strategies may be part of the problem. Sensitivity is inherently lower in crosslink searches than standard proteomic searches because the search space is far larger. However, this reduced search sensitivity can be only part of the problem. Our analysis indicates that large numbers of crosslinked peptides enter the mass spectrometer and are available for selection, but we are simply not triggering enough MS^2^ events to represent the yield that we generate. Interestingly, the total ion chromatogram (TIC) measured after installing the click-precursors looks essentially indistinguishable from the TIC measured after the click reaction (**Figure 5**). It is as if the crosslink signal has vanished.

**Figure 5.**
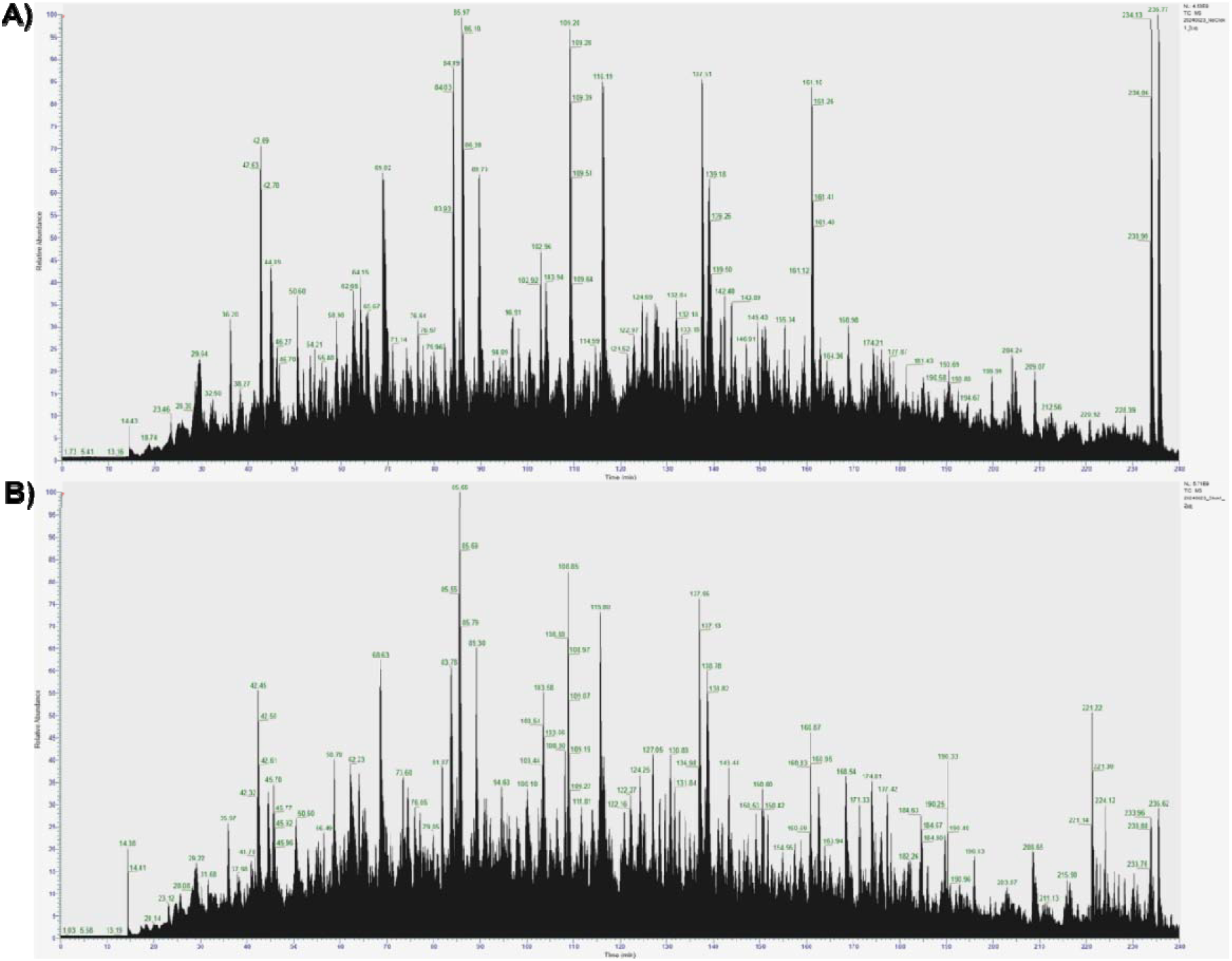
Impact of click-linking on the appearance of the ion population. (A) TIC of a shotgun proteomics analysis of an A549 digest of the labeled, no-click sample. (B) TIC of a shotgun proteomics analysis of an A549 digest of the labeled and clicked sample.

We propose that extensive signal splitting is the reason for poor efficiency in crosslink detection (**Figure 6**). For sparse labeling, where crosslinking networks are local and separated from each other, any surface exposed lysine in a tryptic fragment will distribute its crosslinks across multiple lysine linkage points, and each linkage can be represented by multiple peptides, depending on enzymatic efficiency. These multiple linkages create crosslinked peptides that draw from all peptides, not just the detectable ones, and dilute the signal for any given crosslinked peptides signal across LC retention time and m/z space. For example, a crosslinked peptide comprised of two lysines that are each represented by two peptides, which competes with 5 other crosslinks, can experience a 24-fold dilution compared to a fully non-competitive crosslink. This worsens to 96-fold dilution when including a charge state distribution of 4 peaks for a crosslinked peptide. Of course, the dilution will not be evenly distributed across all 96 channels. There will be a bias based on peptide abundance and the underlying stoichiometry of PPIs, but the dilution is nonetheless considerable. In network theory, a percolation threshold can be reached where networks are all connected^31^. This threshold can be reached when click-linking to levels approaching 30% of total peptide involvement, where the competition for lysines is much worse in this situation. In the example above, a linked lysines can now experience competition from other crosslinks and labeling products, including new categories of reaction products that are only possible in high-yield reactions, such as doubly-crosslinked tryptic peptides and additional missed cleavages. Dilution can approach 2000-fold (**Figure 6**). At the mass spectral level, each dilution leads to signal attrition for a given crosslink and (in the extreme) can wash out the signal completely, at least to levels that do not trigger an MS^2^ acquisition. This conclusion is supported by our observation that the number of measured crosslinks trends to a maximum value as the yield is driven higher (**Figure 2B-D**).

**Figure 6.**
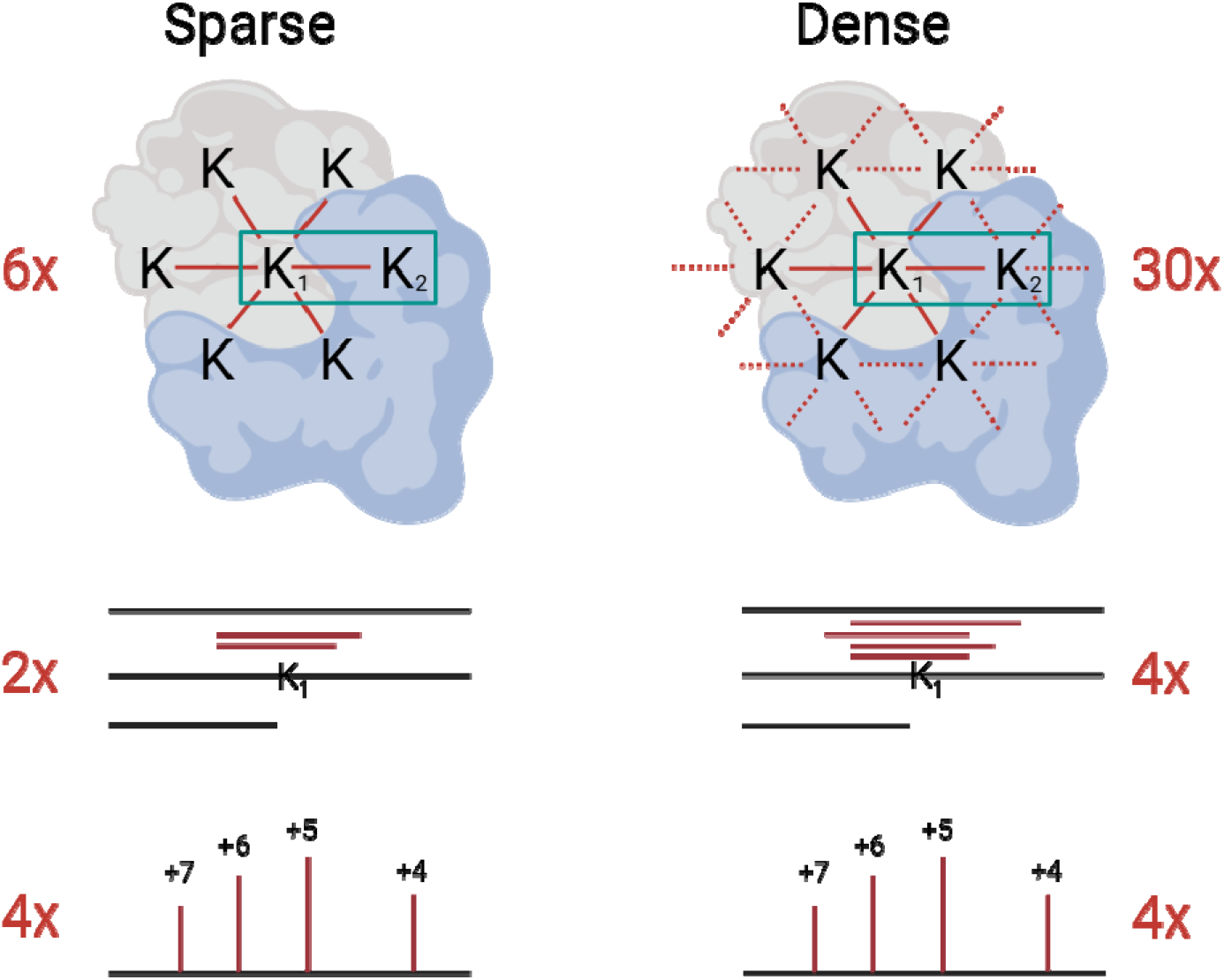
Signal splitting arising from competition for crosslinkable lysines. To estimate signal splitting for a given crosslink (green box) under sparse labeling conditions, where local crosslink networks are isolated, there are 6 competing linkages (top left), ∼2 overlapping peptides for each lysine in the crosslink (middle left), and 4 charge states for each crosslink (bottom left), representing 96 competing MS features for the boxed crosslink. This competition worsens considerably under dense crosslinking conditions, where local networks are linked, leading to 30 competing linkage (top right), ∼4 overlapping peptides for each lysine (due to more missed cleavages resulting from labeling, middle right), 4 charge states (bottom right), representing 1920 competing MS features for the boxed crosslink.

In summary, while the number of crosslinks can increase with higher-yielding chemistry, the signal for any given crosslink will become progressively weaker, at least for lysines that participate in average-sized networks. Stated another way, as the crosslinking yield is driven higher, there are a finite number of lysines that must distribute signal between an exponentially increasing number of crosslinks. At some point the network will saturate, but the number of linkage forms will clearly be orders of magnitude larger than the number of peptides available to represent them. This will lead to significant signal attrition.

There are ways to address the problem, at least in part. Tuning the reaction yield to balance complexity with yield is an obvious approach (and a natural advantage of the click-linking strategy), but it is very likely that each local interaction network will have its own optimal value. Enhancing crosslink detectability with affinity-tagged crosslinkers will reduce ion suppression from linear peptides and increase signal^10^, but it will not help with the complexity of the total pool. Targeted enrichment of crosslinked organelles or tagged proteins should generate the greatest benefit, as these strategies reduce complexity and can be tailored to match the sampling depth of the mass spectrometer. Affinity enrichment and multidimensional separations (including alternative proteases^32^) may always be part of the analytical solution, but other improvements can also improve performance. For example, charge enrichment techniques such as ion mobility^33^ and multiple charge separation technology^34^ would enhance S/N for the higher charge state peptides, and there is room for MS^2^ methods that are better at high charge state selection. All of these strategies will help, but they may not fully overcome the challenge of high sample complexity in the fully-crosslinked proteome, a detection problem that is much worse than standard bottom-up proteomics.

## Data Availability

The crosslinking and labeling data generated in this study have been deposited in the PRIDE partner repository^35^ with the dataset identifier PXD069512.

## Supporting Information

Figure S1: Investigation of click-link side reactions. Figure S2: Lysine labeling yield and distribution of a 1 mM reagent installation reaction. Figure S3: Reduction in crosslink precursor intensity upon click-linking. Figure S4: Example MS^2^ spectra of click-linked peptides. Figure S5: Crosslink spectrum matches as a function of database search tool. Table S1: Crosslinked spectrum matches from alternative database search engines.

## Author contributions

Conceptualization, D.C.S.; methodology and experiment design, D.C.S., N.I.B, B.C.A., A.R.M.M.; experiments, N.I.B., B.C.A. P.D. and M.F.K.; project organization, resources and funding acquisition. D.C.S.; data analysis and visualization, B.C.A, N.I.B. and D.A.C.; manuscript writing, D.C.S and B.C.A.

## Acknowledgements

This work was funded by the Natural Sciences and Engineering Research Council of Canada Discovery Grants RGPIN 2017-04879 to D.C.S. All figures created and/or assembled with BioRender.com, released under a Creative Commons Attribution-NonCommercial-NoDerivs 4.0 International license.

## Notes

The authors declare no competing financial interest.

