## Supporting Information for "Signal Attrition in Whole Cell Crosslinking Mass Spectrometry"

Dr. David Schriemer

Department of Biochemistry and Molecular Biology, Cumming School of Medicine

University of Calgary, 3330 Hospital Drive NW T2N-4N1

____________________________________________________________________________________

^#^These authors contributed equally to the study.

**Table of Contents:**

1. **Figure S1.** Investigation of click-link side reactions.
2. **Figure S2.** Lysine labeling yield and distribution of a 1 mM reagent installation reaction.
3. **Figure S3.** Reduction in crosslink precursor intensity upon click-linking.
4. **Figure S4.** Example MS^2^ spectra of click-linked peptides
5. **Figure S5.** Crosslink spectrum matches as a function of database search tool.
6. **Table S1.** Distribution of crosslink products under different digestion protocols.
7. **Table S2.** Crosslinked spectrum matches from alternative database search engines.

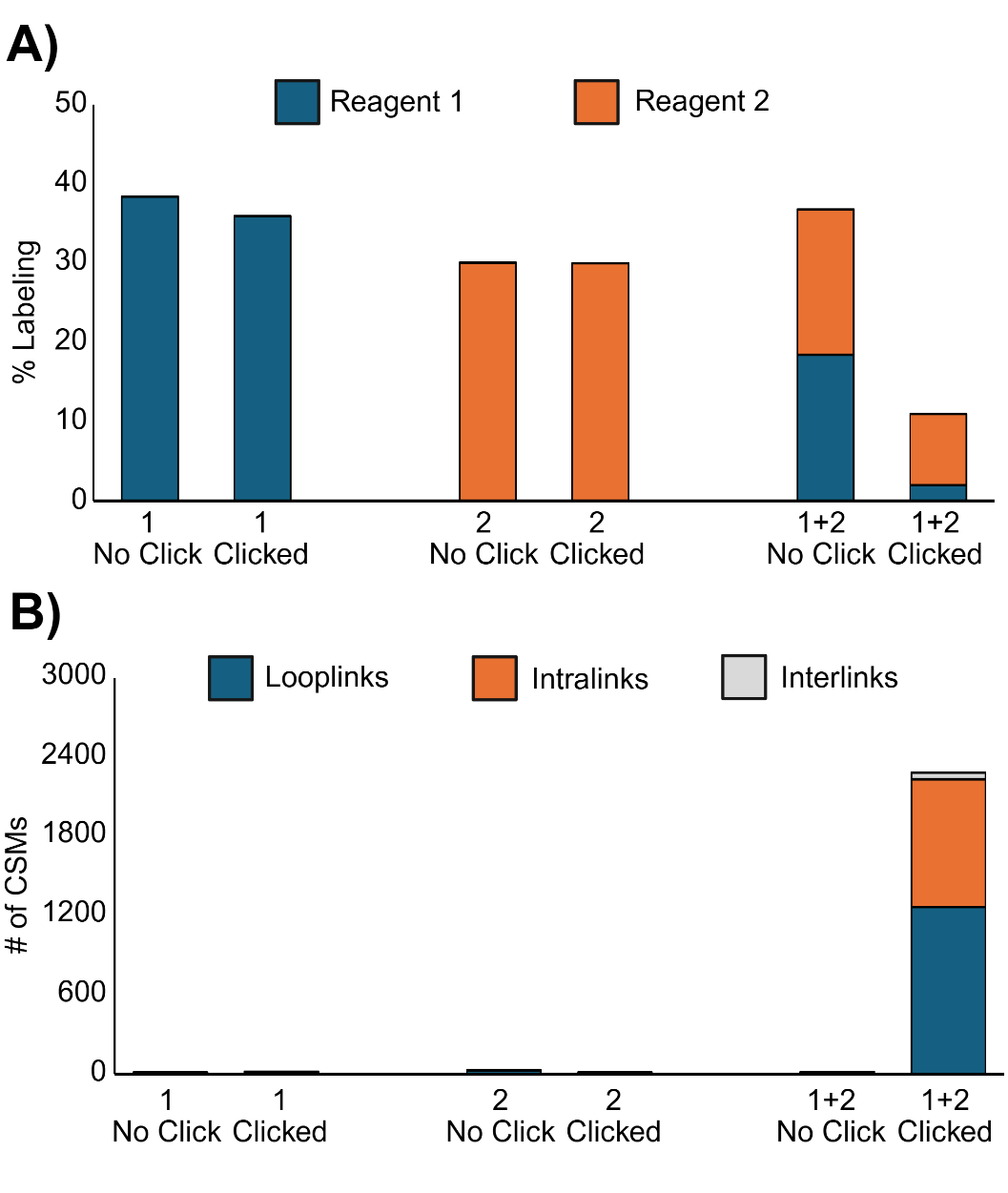

**Figure S1.** Inspection of possible side products resulting from click-linking of A549 cells. (A) Levels of precursor peptides detected via shotgun analysis before and after click-linking, under three different conditions: only installation of 1 mM reagent **1** (the alkyne, left panel), only installation of 1 mM reagent **2** (the azide, middle panel) and in the presence of both reagents (1 mM each). The data indicate no appreciable reduction in precursors arising other than through the intended reaction products (protein crosslinks). (B) as in A, but showing crosslink spectrum matches (CSMs).

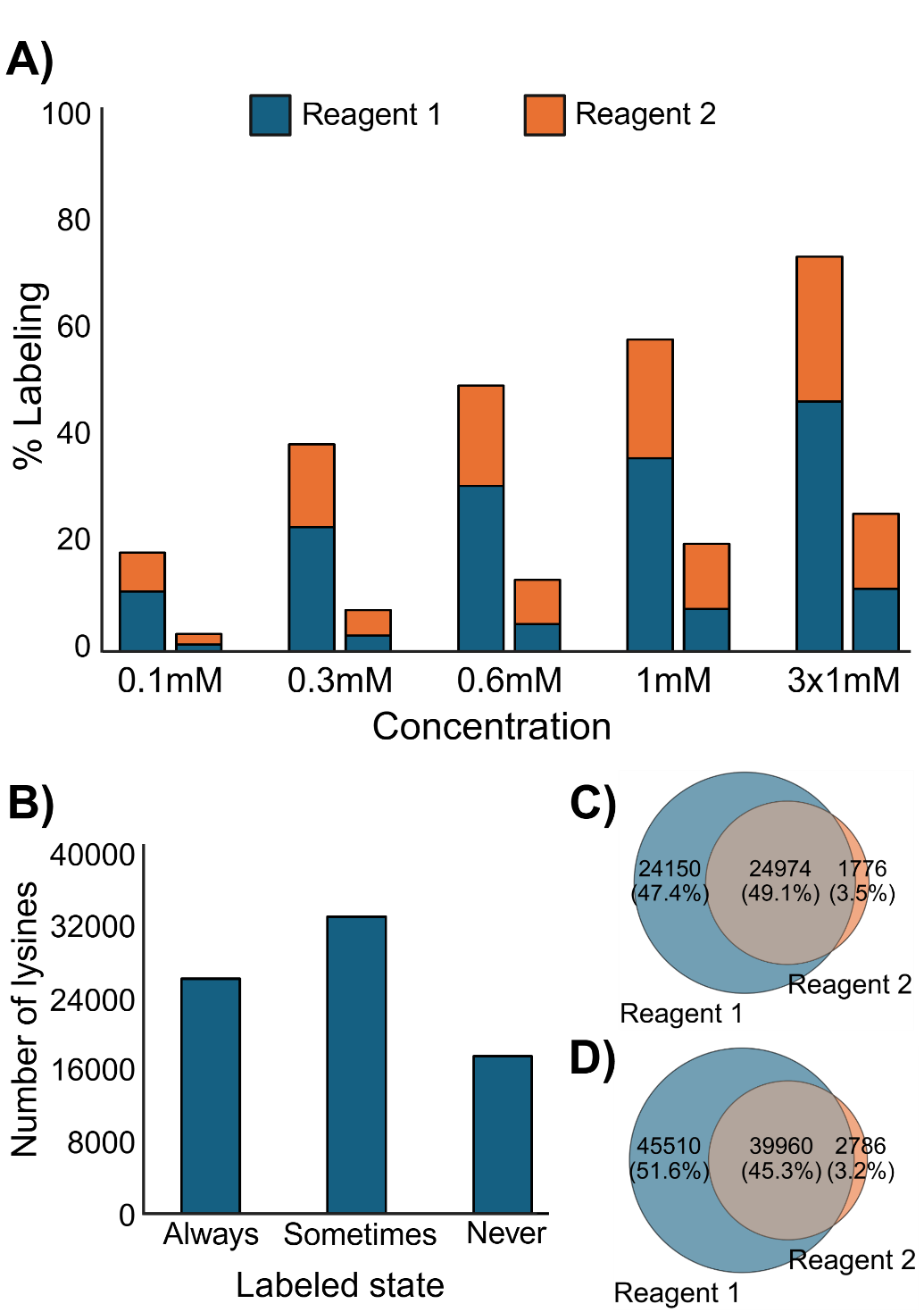

**Figure S2.** Lysine labeling yield and distribution of a 1 mM reagent installation reaction and clicking. (A) yield of reagents 1 and 2 installation before and after clicking considering only lysine-containing peptides in the proteome; (B) distribution of labeling states for *unique* lysines in a full proteome sample. Each bar represents the union between reagent 1 and 2; (C) distribution of labeled unique lysines between reagent 1 and 2, from the Sometimes and Always sets in (B); (D) same as (C), but at the unique peptide level instead of unique lysine.

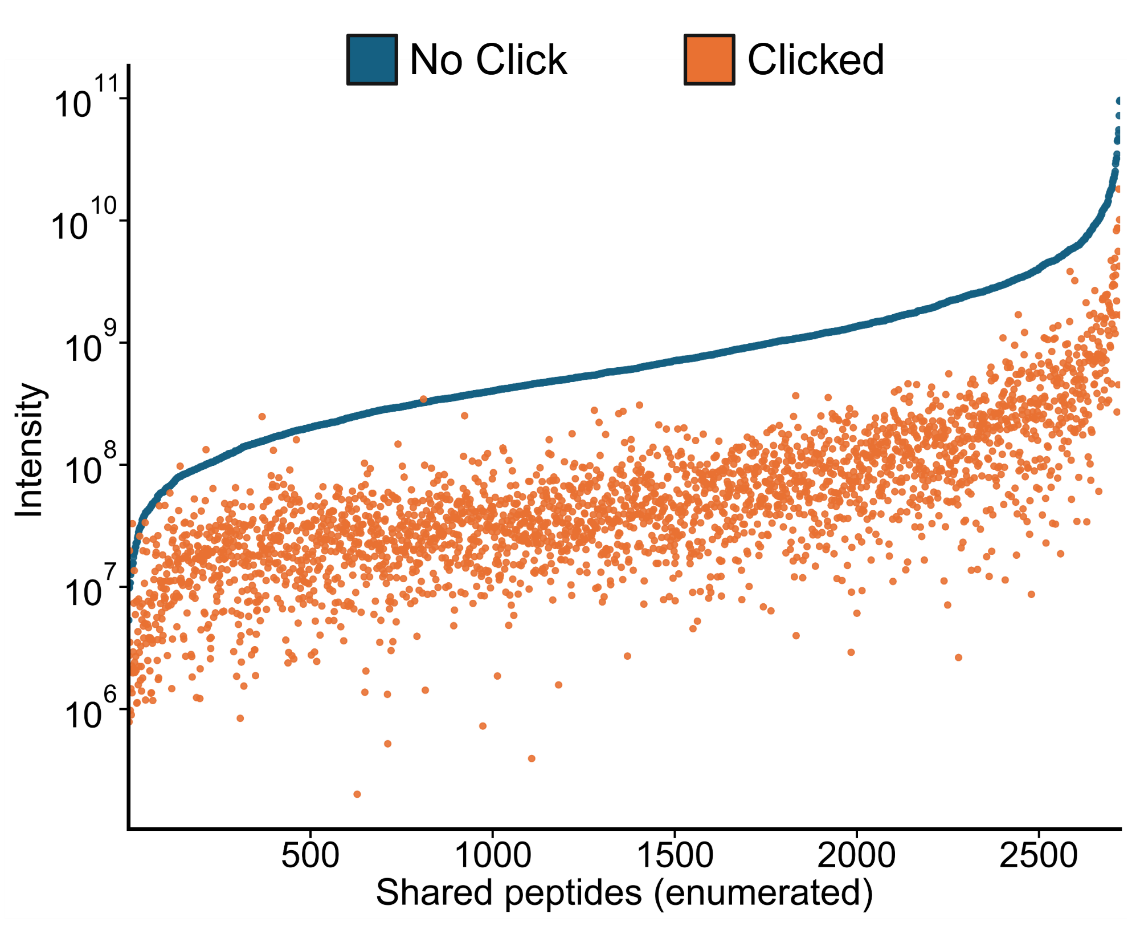

**Figure S3.** Reduction in peptide precursor intensity upon click-linking.

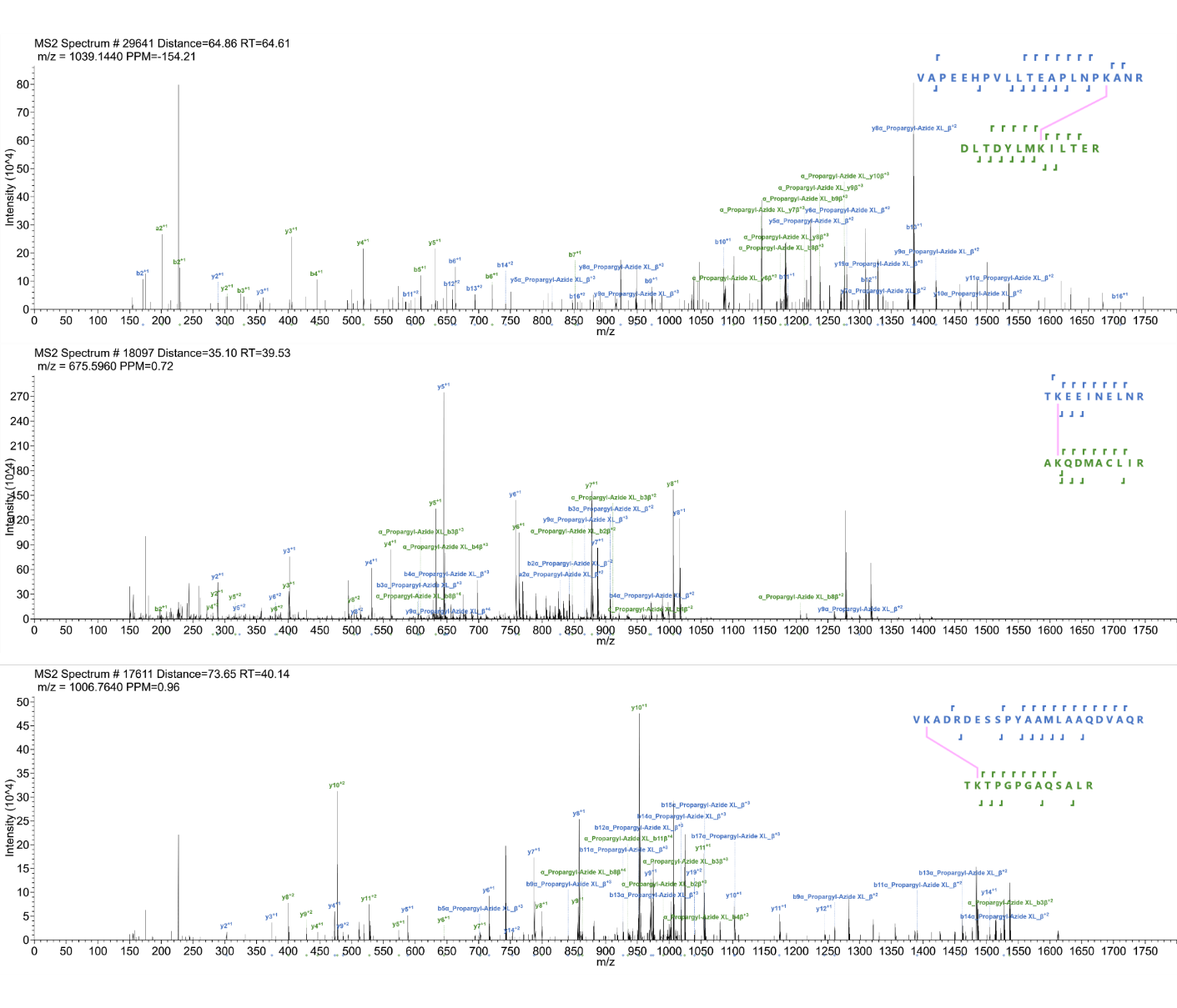

**Figure S4**. Example MS^2^ spectra of click-linked peptides. No evidence of triazole cleavage is found, although some minor amide cleavage at the ε nitrogen in lysine is seen, as is typical with NHS-based linkages.

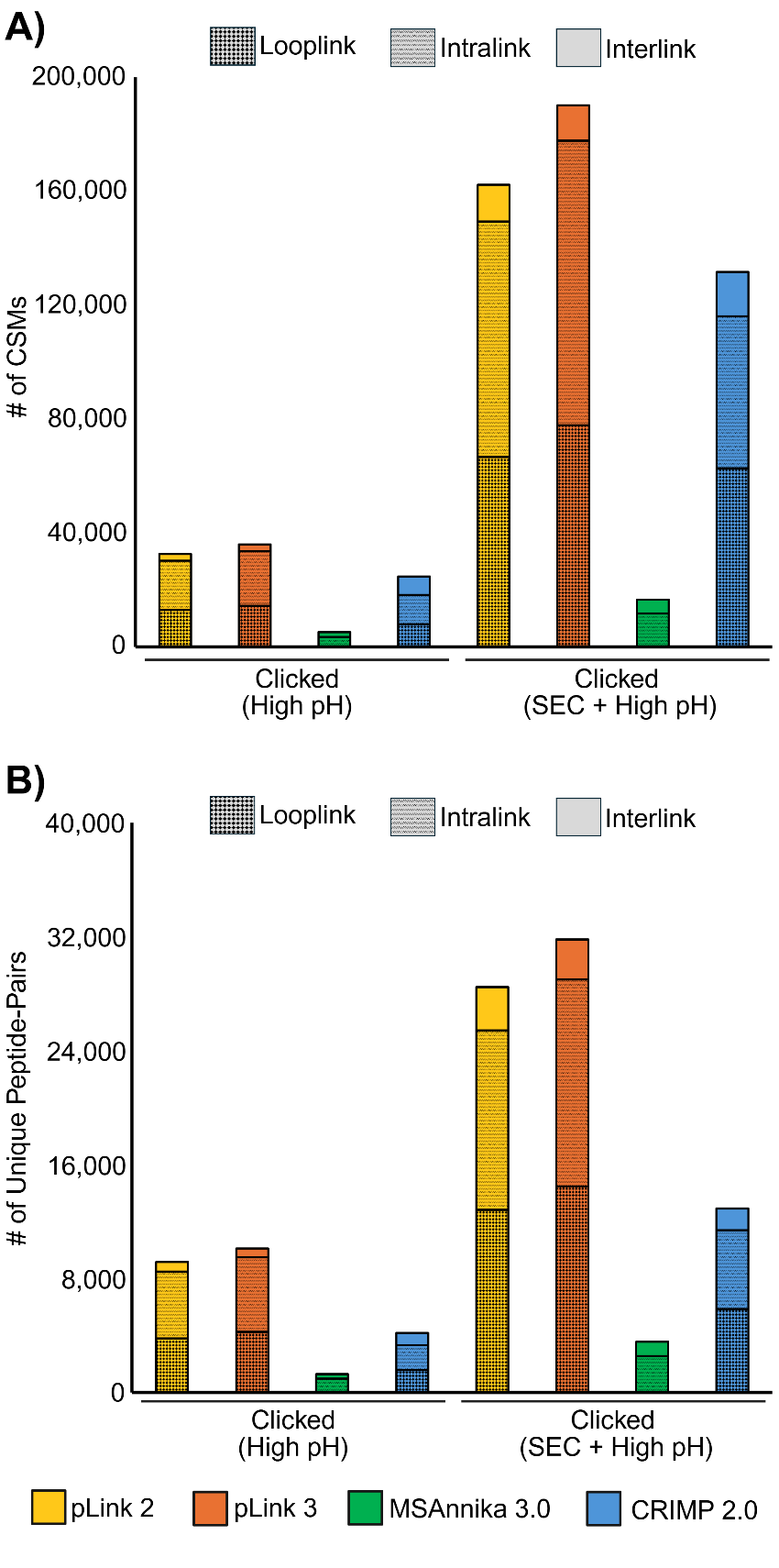

**Figure S5.** Crosslink Spectrum Matches and Unique Peptide-Pairs as a function of database search tool for Click-Linking samples. MS Annika 3.0 does not report looplink matches in its results.
